# Loss of cilia drives centriole clustering and elimination during mammalian spermatogenesis

**DOI:** 10.64898/2025.12.04.692363

**Authors:** Jun Jie Chen, Xiangyu Gong, Michael Mak, Feng-Qian Li, Ken-Ichi Takemaru

## Abstract

Cilia are microtubule-based organelles essential for signaling and motility, and their dysfunction causes ciliopathies often associated with infertility. In male germ cells, two types of cilia are present: zygotene primary cilia and sperm flagella. To define the role of cilia in spermatogenesis, we conditionally ablated the distal appendage protein CEP164, required for basal body docking and ciliogenesis, in male germ cells. CEP164 localized to the mother centriole/basal body throughout spermatogenesis, and its loss led to male infertility accompanied by absence of both zygotene cilia and sperm flagella. Despite defective ciliogenesis, meiotic chromosome pairing and DNA double-strand break repair proceeded normally. However, round spermatids exhibited basal body docking and flagellogenesis defects, and frequently formed supernumerary centriole clusters that were subsequently eliminated via residual bodies. Live-cell imaging revealed that centrioles were highly mobile, and centriole pairs from neighboring cells often associated through intercellular bridges, forming aggregates. These results establish that basal body docking is crucial for retaining centrioles within spermatids, and its disruption leads to centriole clustering and loss. In contrast, zygotene cilia are dispensable for meiotic chromosome pairing and DNA repair during mammalian spermatogenesis.

## Introduction

Cilia are highly conserved, microtubule-based organelles that extend from the surface of many different cell types (Hilgendorf *et al*, 2024; Hyland & Brody, 2021; Mill *et al*, 2023; Moran *et al*, 2024; Wallmeier *et al*, 2020). Based on their axonemal architecture, cilia are broadly classified into two types: non-motile primary cilia, with a 9+0 microtubule arrangement, and motile multicilia, with a 9+2 arrangement. Primary cilia are ubiquitous and function as sensory organelles that respond to chemical and mechanical cues in the environment. In contrast, multicilia are restricted to specialized epithelia, such as those of the airways, reproductive tracts, and the brain ventricles, where they generate directional fluid flow through coordinated beating. Sperm flagella represent a unique variant: singular motile organelles with a 9+2 axoneme and elaborate accessary structures that are indispensable for motility. Dysregulation of cilia causes a spectrum of multisystemic disorders known as ciliopathies, which frequently include male infertility. To date, mutations in at least 247 genes have been associated with ciliopathies, affecting either or both ciliary types (Mill *et al*., 2023; Moran *et al*., 2024).

Ciliogenesis is initiated during the G0/G1 phase of the cell cycle, when the mother centriole transforms into a basal body that first engages with small vesicles or the plasma membrane prior to axoneme assembly (Hilgendorf *et al*., 2024; Mill *et al*., 2023). This direct basal body-membrane docking is mediated by distal appendages, pinwheel-like fibrous structures emanating from each microtubule triplet (Kanie *et al*, 2025; Ma *et al*, 2023). Distal appendages are composed of several core proteins, including centrosomal protein of 164 kDa (CEP164). CEP164 is critical for small vesicle recruitment and basal body docking during both primary and multiciliogenesis (Schmidt *et al*, 2012; Siller *et al*, 2017). CEP164 recruits downstream ciliary proteins, including Cby1 and ciBAR family members, to promote basal body docking and ciliogenesis (Burke *et al*, 2014; Hoque *et al*, 2021; Li *et al*, 2016; Siller *et al*., 2017). We previously demonstrated that multiciliated cell-specific CEP164-knockout (KO) mice exhibit a profound loss of multicilia in the trachea, oviduct, and ependyma, with ∼20% succumbing to severe hydrocephalus (Siller *et al*., 2017). More recently, we reported that male mice are infertile due to dysfunctional multicilia in the efferent ducts connecting the testis and epididymis (Hoque *et al*., 2021). In striking contrast to its essential role in mammalian ciliogenesis, the *Drosophila* CEP164 ortholog is dispensable for basal body docking, ciliogenesis, and male fertility (Hou *et al*, 2023). However, the role of CEP164 in mammalian spermatogenesis has remained unknown. Beyond its ciliary functions, CEP164 has also been implicated in DNA damage responses (Chaki *et al*, 2012; Pan & Lee, 2009), although this role has been challenged (Daly *et al*, 2016).

Mammalian spermatogenesis is a complex and tightly regulated process that generates haploid gametes (spermatozoa) from diploid spermatogonial stem cells within the seminiferous tubules of the testis (Griswold, 2016; Han, 2024; Teves & Roldan, 2022). In mice, the process is organized into 12 stages and completed in approximately 34.5 days (Oakberg, 1956). Spermatogenesis can be broadly categorized into three major phases: (1) self-renewal of spermatogonial stem cells, 2) meiosis, in which spermatocytes give rise to spermatids, and 3) spermiogenesis, the differentiation of spermatids into mature spermatozoa, including flagellogenesis. A unique feature of both male and female gametogenesis is the formation of stable intercellular bridges that physically interconnect developing germ cells, creating a syncytial network (Gerhold *et al*, 2022; Greenbaum *et al*, 2011). The diameter of these cytoplasmic bridges can reach up to 3.0 μm, allowing the exchange of RNAs, proteins, and organelles, thereby ensuring their synchronized development and functional coordination within the cyst (Greenbaum *et al*., 2011).

Recent studies in zebrafish have reported the presence of primary cilia in both primary oocytes and spermatocytes (Mytlis *et al*, 2022; Xie *et al*, 2022). In oocytes, primary cilia are detected at leptotene and zygotene stages and are essential for proper formation of the meiotic chromosomal bouquet, an evolutionarily conserved configuration that facilitates synaptonemal complex assembly and homologous chromosome pairing (Mytlis *et al*., 2022; Mytlis *et al*, 2023). During bouquet formation, telomeres slide along perinuclear microtubules and cluster at a small region of the inner nuclear membrane, creating the “floral arrangement”, hence the name. Strikingly, loss of zygotene cilia disrupts synaptonemal complex formation, impairs oogenesis, and leads to infertility.

However, this conclusion is in contrast to previous findings indicating that germ cell-specific removal of cilia exhibits no significantly effects on oogenesis and fertility in zebrafish females, while causing infertility in males (Borovina & Ciruna, 2013; Liu *et al*, 2023; Xie *et al*., 2022). In zebrafish spermatocytes, primary cilia are observed from leptotene through diplotene stages (Xie *et al*., 2022). Germ cell-specific ablation of primary cilia results in defective DNA double-strand break repair (DSB), increased apoptosis, and infertility in males. In mice, primary cilia are detected at the zygotene stage in the adult testis (Lopez-Jimenez *et al*, 2022). Notably, only about 20% of zygotene spermatocytes extend primary cilia. Analysis of the timing of ciliogenesis relative to bouquet formation indicated that primary cilia are not directly involved in bouquet formation, suggesting that zygotene cilia are dispensable for homologous chromosome pairing in mammalian spermatocytes. Nevertheless, the role of primary cilia in mammalian germ cells remains to be elucidated.

To investigate the role of CEP164 and primary cilia in mammalian spermatogenesis, we generated male germ cell-specific *Cep164*-KO mice. We found that cilia were largely absent in zygotene spermatocytes and sperm, and males were infertile. Notably, meiotic chromosome pairing and DSB repair progressed normally in the absence of primary cilia. In contrast, loss of CEP164 caused impaired basal body docking and defective flagellogenesis in spermatids. Unexpectedly, undocked centrioles were highly mobile and associate with one another likely through intercellular bridges to form aggregates, most of which were ultimately discarded into residual bodies in the lumen of seminiferous tubules. In sum, our findings demonstrate that basal body docking mediated by CEP164 is essential for centriole immobilization and retention, and flagellogenesis, whereas zygotene cilia are dispensable for meiotic chromosome pairing or DNA repair during mammalian spermatogenesis.

## Results

### Loss of CEP164 in male germ cells causes infertility

To investigate the biological role of cilia in spermatogenesis, we conditionally deleted *Cep164* in male germ cells by crossing *Cep164^fl/fl^* mice (Hoque *et al*., 2021; Siller *et al*., 2017) with *Stra8*-iCre, which transiently expresses Cre recombinase from type A spermatogonia to preleptotene spermatocytes (Sadate-Ngatchou *et al*, 2008).

CEP164 consists of 1,460 amino acids and contains a WW domain and three coiled-coil domains (Fig EV1A). Cre-mediated recombination excises exon 4, resulting in a frameshift and premature truncation at amino acid 65 (Fig EV1B). These mice are hereafter referred to as cKO. Genotypes were confirmed by PCR analysis of tail DNA (Fig EV1C). Immunofluorescence (IF) staining showed that CEP164 was largely absent from the mother centriole and ciliary base in cKO testes (Fig EV1D).

To examine the biological importance of CEP164 during spermatogenesis, we evaluated male fertility by mating *Cep164^fl/fl^* control and cKO males with *Cep164^fl/fl^* control females. cKO males failed to produce any offspring (Table 1), indicating infertility. To shed light on the underlying cause, we examined testis morphology, weight, and sperm production. Testes from cKO mice appeared grossly normal (Fig 1A), and testis weights were comparable to those of controls (Fig 1B). However, cKO mice exhibited a dramatic reduction in the number of mature sperm harvested from the cauda epididymis (Fig 1C). Consistent with this finding, hematoxylin and eosin (H&E) staining revealed an absence of sperm in the epididymal lumen of cKO males (Fig 1D). These results demonstrate that CEP164 is essential for male fertility in mice.

**Figure 1.**
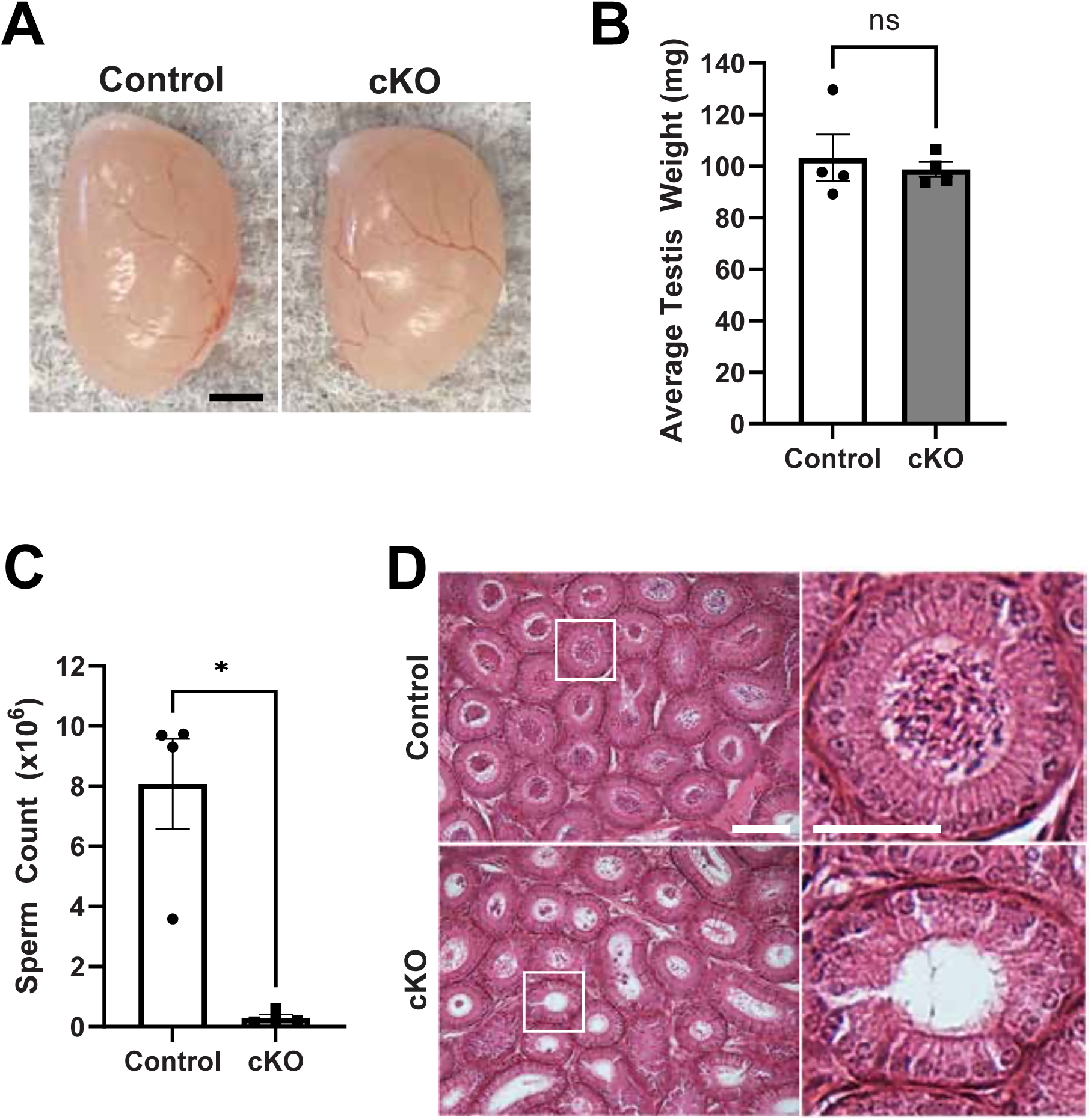
*Stra8-iCre;Cep164^fl/fl^* males are infertile with reduced sperm counts. (A) Representative images of adult testes from *Cep164^fl/fl^* (control) and *Stra8-iCre;Cep164^fl/fl^* (cKO) mice. Scale bar, 2 mm. Quantification of testis weight (B) and sperm count (C). Adult mice (2-5-month-old) were euthanized, and testes were collected for weight measurements. Sperm were isolated from the cauda epididymis and counted using a hemocytometer. n = 4 per genotype. Error bars represent ± SEM. Welch’s t test: not significant (ns), *P < 0.05. (D) Representative images showing H&E staining of adult epididymal sections. Scale bars, 100 and 50 µm (enlarged images).

**Table 1.**
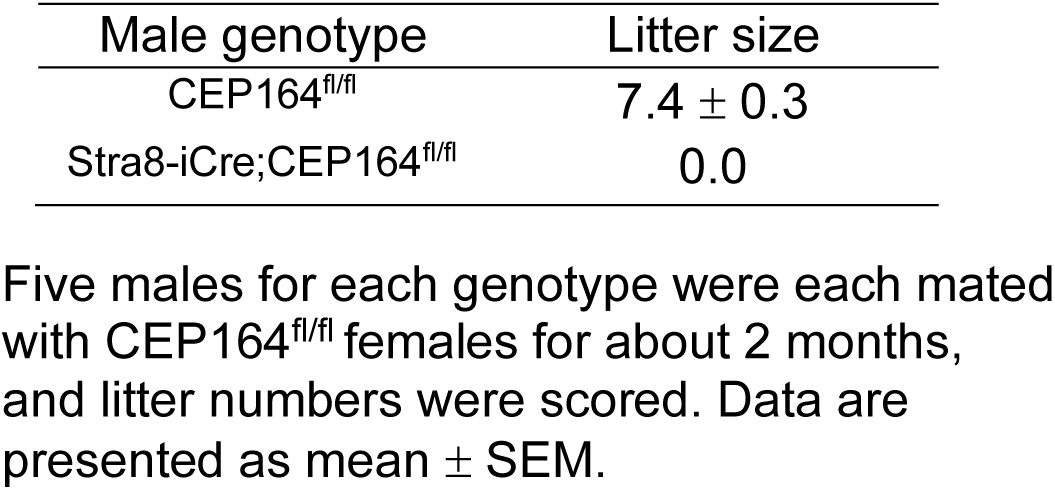
Fertility test of Stra8-Cre;CEP164^fl/fl^ male mice.

| Male genotype | Litter size |
| --- | --- |
| CEP164<sup>fl/fl</sup> | 7.4 ± 0.3 |
| Stra8-iCre;CEP164<sup>fl/fl</sup> | 0.0 |

Five males for each genotype were each mated with CEP164<sup>fl/fl</sup> females for about 2 months, and litter numbers were scored. Data are presented as mean ± SEM.

### Defective flagellogenesis and aberrant sperm head elongation in cKO testes

To gain insight into spermatogenesis defects in cKO mice, we performed periodic acid-Schiff (PAS) staining on adult testis sections. In *Cep164^fl/fl^* control sections, elongated sperm flagella were readily observed in the lumen (Fig 2A, asterisk). In marked contrast, cKO testes lacked detectable flagella. In addition, we observed abnormally elongated sperm heads in cKO sections (Fig 2A, arrowheads). This was more clearly visualized by co-staining with lectin peanut agglutinin (PNA; acrosome marker) and DAPI (Fig 2B, arrowheads). IF staining for acetylated α-tubulin (A-tub; a marker of flagella and stable microtubules) further confirmed the absence of flagella in cKO seminiferous tubules (Fig 2C, asterisk). Notably, the manchette, a transient microtubule structure required for sperm head shaping, appeared to be abnormally elongated and persisted in cKO spermatids (Fig 2C, arrowheads), likely contributing to the defective head morphology. Similar elongated head structures have been reported in mouse models of ciliary defects (Lehti *et al*, 2013; San Agustin *et al*, 2015; Zhang *et al*, 2016). Taken together, these findings demonstrate that, in contrast to *Drosophila* spermatogenesis, CEP164 is essential for mammalian sperm flagellogenesis.

**Figure 2.**
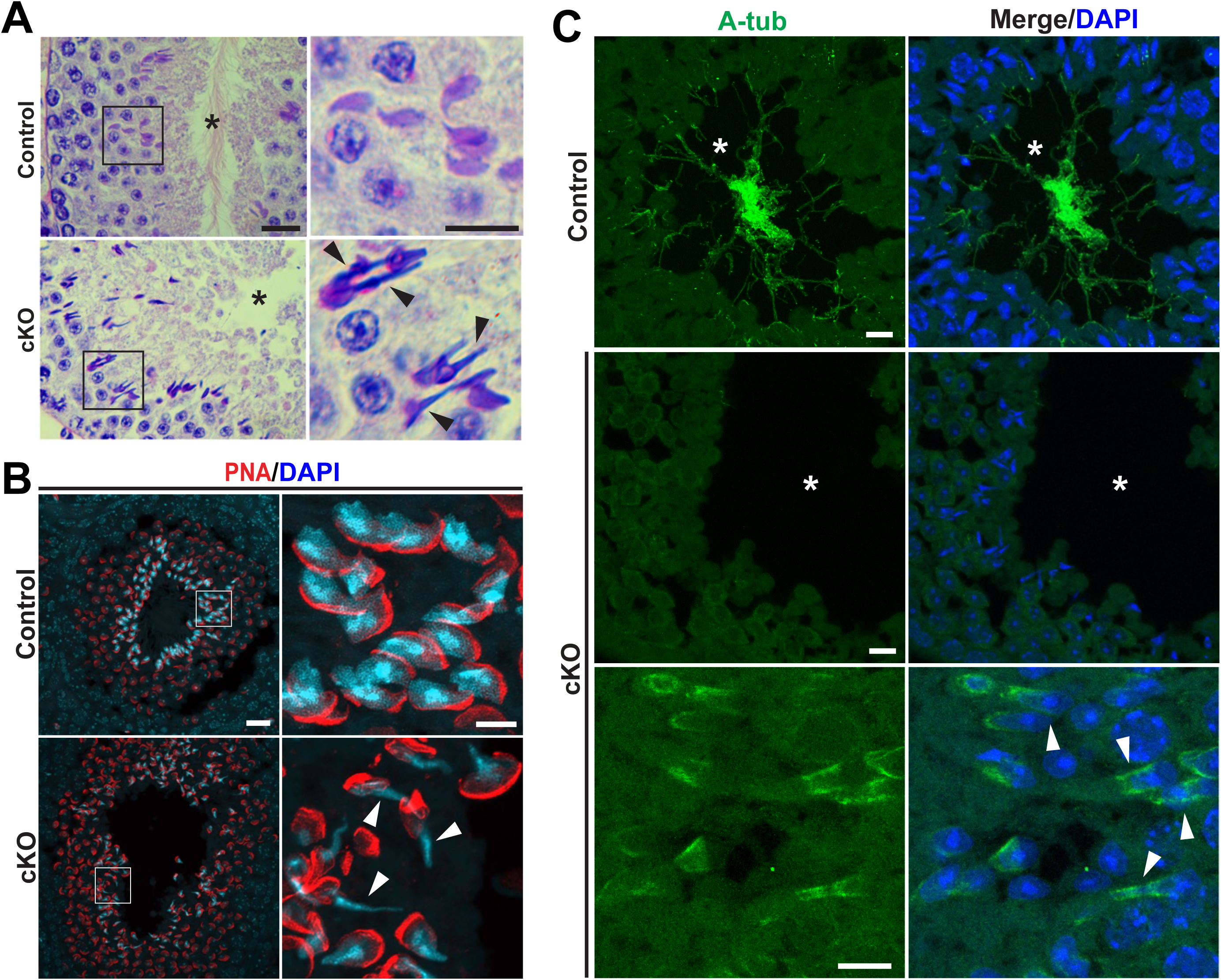
Histological and immunofluorescence analysis of testes. (A) Periodic acid-Schiff (PAS) staining of seminiferous tubules from adult control and cKO mice. Asterisks indicate the lumen where sperm flagella are typically observed. Arrowheads mark abnormally elongated sperm heads in cKO tubules. Scale bars, 20 and 10 µm (enlarged images). (B) Paraffin sections of adult testes stained with lectin peanut agglutinin (PNA; acrosome marker) and DAPI (nuclei). Arrowheads highlight abnormally elongated sperm heads in cKO tubules. Scale bars, 20 and 5 µm (enlarged images). (C) IF staining of adult paraffin testis sections with acetylated α-tubulin (A-tub) antibody (marker of flagella and stable microtubules). Asterisks denote the lumen. Arrowheads point to abnormally elongated sperm heads in cKO tubules with persistent manchettes. Scale bars, 10 µm.

### Depletion of zygotene cilia does not significantly impair meiotic chromosome pairing or DNA repair during murine spermatogenesis

In zebrafish oogenesis, primary cilia in zygotene spermatocytes are required for meiotic bouquet formation and subsequent homologous chromosome pairing (Mytlis *et al*., 2022). Our cKO mouse model provided us an opportunity to investigate the role of zygotene cilia in spermatogenesis. To facilitate the visualization of centrioles, we utilized the ROSA-EGFP-Centrin1 mouse line, which ubiquitously expresses EGFP-Centrin1 in all cell types (Hirai *et al*, 2016).

To quantify the proportion of zygotene spermatocytes bearing primary cilia, we prepared seminiferous tubule squashes from adult testes, followed by IF staining for A-tub and synaptonemal complex 3 (SYCP3; a meiotic chromosomal axis marker) (Fig 3A). In control testes, primary cilia, typically long (15-25 μm) and emanating from centrioles (Fig 3A, arrowheads), were detected in 18.5% of zygotene spermatocytes (Fig 3B), in agreement with previous observations (Lopez-Jimenez *et al*., 2022). In marked contrast, no primary cilia were observed in cKO zygotene spermatocytes.

**Figure 3.**
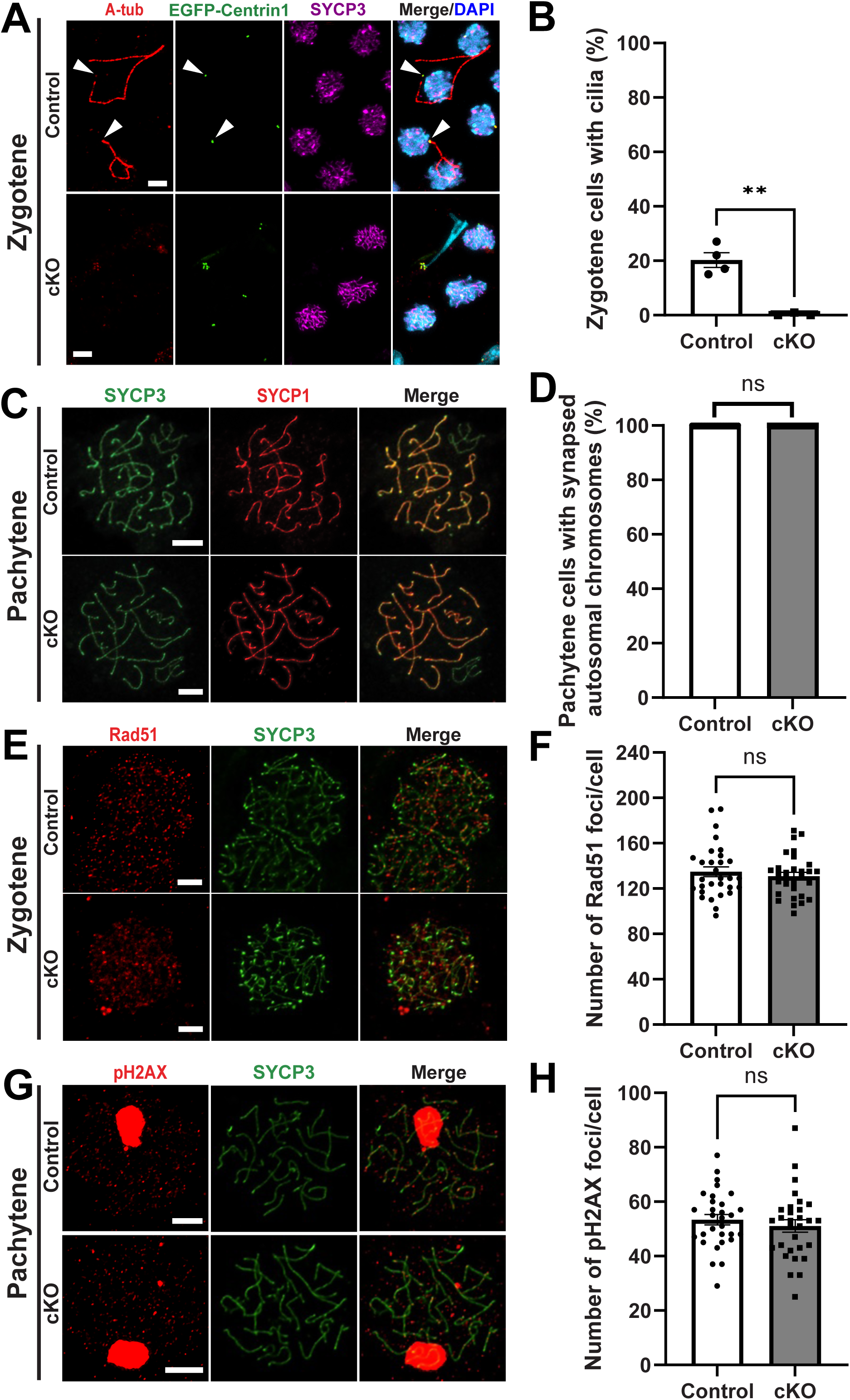
Loss of zygotene cilia does not overtly affect meiotic chromosome pairing or DNA repair. (A) IF staining of seminiferous tubule squashes from control or cKO mice expressing EGFP-Centrin1 (centriole marker) with antibodies against A-tub and SYCP3 (meiotic chromosomal axis marker). Arrowheads indicate basal bodies extending primary cilia in zygotene cells. Scale bar, 5 µm. (B) Quantification of zygotene cilia. n = 4 control and n = 3 cKO mice, with ∼100 zygotene cells scored per mouse. Error bars represent ± SEM. Welch’s t test: **P < 0.01. (C) IF staining of chromosome spread preparations for SYCP3 and SYCP1 (marker of synapsed regions of the synaptonemal complex). Scale bar, 5 µm. (D) Quantification of pachytene spermatocytes with fully synapsed autosomal chromosomes. n = 3 per genotype, with 10 cells analyzed per mouse. Welch’s t test: not significant (ns). (E, G) IF staining of chromosome spread preparations of zygotene spermatocytes (E) for SYCP3 and Rad51 (DNA recombinase mediating homologous strand exchange) and (G) pachytene spermatocytes for SYCP3 and pH2AX (marker of DNA double-strand break sites). Scale bars, 5 µm. (F, H) Quantification of foci for each DNA repair marker. n = 3 per genotype, with 10 cells quantified per mouse. Error bars represent ± SEM. Welch’s t test: not significant (ns).

Next, we evaluated whether loss of cilia influences meiotic chromosome pairing in cKO spermatocytes. Chromosome spread slides were prepared from dissociated seminiferous tubules and immunostained for SYCP1 and SYCP3 (a marker of synapsed regions of the synaptonemal complex) (Fig 3C). SYCP3 decorates the lateral elements of both paired and unpaired chromosomal regions, whereas SYCP1 localizes to the central elements of paired chromosomes. We observed that all 19 autosomes were fully paired in both control and cKO pachytene spermatocytes, indicating that homologues synapsis occurs normally without primary cilia (Fig 3D).

During prophase I, homologous chromosome pairing is closely coordinated with DNA double-strand break (DSB) formation and repair. CEP164 has been implicated in DNA damage response pathways (Chaki *et al*., 2012; Sivasubramaniam *et al*, 2008), and in zebrafish, primary cilia play a critical role in meiotic homologous recombination repair in spermatocytes (Xie *et al*., 2022). To assess if cKO spermatocytes show defects in DSB repair processes, we performed IF staining for key DNA repair proteins: Rad51, a critical recombinase mediating homologous strand exchange, and phospho-histone H2A.X (γH2AX [Ser139]), which marks DSB sites. The number of foci for these markers was comparable between control and cKO spermatocytes (Fig 3E-H), indicating that DSB repair was not significantly impaired. Thus, these results suggest that primary cilia, unlike in zebrafish, are not essential for meiotic chromosome pairing and DNA repair in the mouse testis.

### Basal body docking defects and centriole clustering in cKO spermatids

During the course of our study, we unexpectedly observed abnormal aggregation of multiple centrioles in cKO testes (Fig 4A, yellow arrowheads). To ensure detection accuracy, centrioles were visualized using two independent centrin antibodies, Centrin 20H5 and Centrin 3, as indicated. Whereas each spermatid typically contains a single pair of centrioles (Fig 4A, white arrowheads), strikingly, large clusters comprising as many as 23 centrioles per cell were detected in cKO testes (Fig 4B). In the cKO testis, centriole clusters were evident as early as postnatal day 22 (P22) (Fig 4C, arrowhead), shortly after the appearance of round spermatids during the synchronized first wave of spermatogenesis (Bellvé *et al*, 1977; Ernst *et al*, 2019). Analysis by transmission electron microscopy (TEM) revealed basal body docking defects and the presence of at least three centrioles in cKO step 8 spermatids (Fig 4D). Notably, these centrioles failed to dock to the nuclear envelope but developed a structure resembling the basal plate (Fig 4D, arrowhead), which is part of the head-tail coupling apparatus (HTCA) that normally forms between the proximal centriole and nuclear membrane (Buglak *et al*, 2025; Wu *et al*, 2020). To assess whether centriole clusters in cKO testes preserve the normal 1:1 ratio of mother to daughter centrioles, we performed IF staining of adult testis sections using antibodies against centrin and ODF2 (a mother centriole marker). As shown in Fig 4E, centriole clusters in cKO testes retained the expected mother-to-daughter ratio. Together, these data suggest that loss of CEP164 results in basal body docking defects and abnormal centriole aggregation in spermatids.

**Figure 4.**
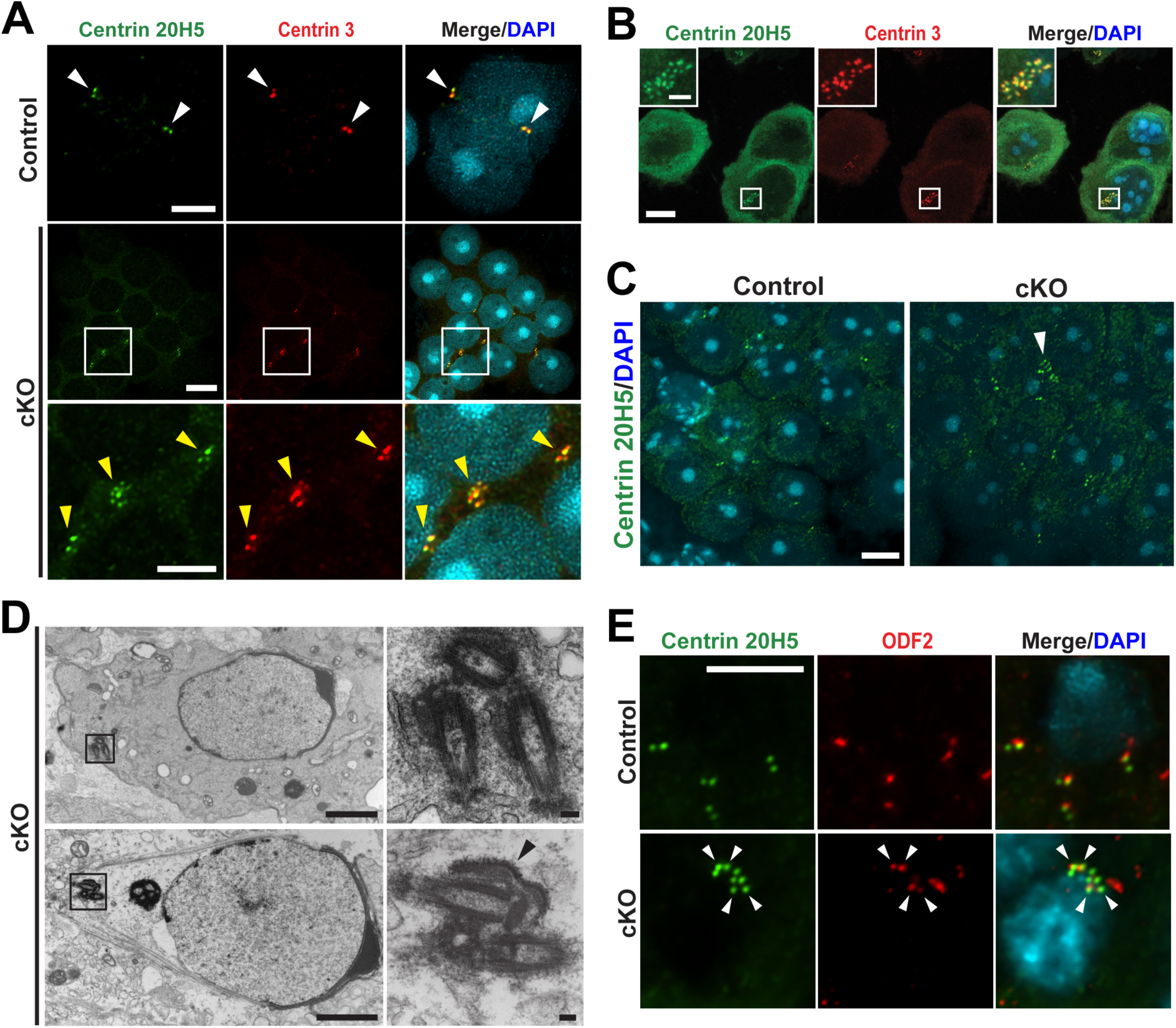
Abnormal clustering of supernumerary centrioles in cKO spermatids. (A, B) IF staining of chromosome spreads from cKO testes using Centrin 20H5 and Centrin 3 antibodies. (A) Enlarged views of boxed regions are shown below. In control round spermatids, two centrioles are typically present (white arrowheads). In contrast, cKO round spermatids exhibit abnormal aggregates of multiple centrioles (yellow arrowheads). Scale bars, 10 and 5 µm (enlarged images). (B) Representative example showing 23 centrioles within a single cKO round spermatid. Scale bars, 20 µm and 5 µm (insets). (C) IF staining of P22 paraffin testis sections using Centrin 20H5 antibody. The arrowhead indicates a centriole cluster. Scale bar, 5 µm. (D) TEM images of cKO testis sections showing the presence of three undocked centrioles in a single step 8 spermatid. The arrowhead denotes basal plate-like structures typically present at the head-tail junction. Enlarged views of boxed regions are shown to the right. Scale bars, 2 µm and 100 nm (enlarged images). (E) IF staining of frozen testis sections using Centrin 20H5 and ODF2 (mother centriole marker) antibodies. Arrowheads point to mother centrioles. Scale bar, 10 µm.

### Altered localization of sperm tail components in cKO elongated spermatids

We next investigated the localization of structural components of the sperm tail. IF staining of adult testis sections were performed using antibodies against centrin and specific sperm tail markers. Sept4 is a major component of the annulus that is present as a distinct ring at the base of flagellum distal to the centrin-positive basal body during spermiogenesis steps 9-14 (Hoque *et al*, 2024; Whitfield, 2024). Whereas such Sept4 rings were readily detectable at the flagellar base in controls, no clear Sept4 rings were observed in cKO spermatids (Fig 5A). Instead, elongated Sept4-positive fibrous structures were frequently noted. ODF2, an outer dense fiber protein of the midpiece, and AKAP3, a fibrous sheath component of the principal piece, also displayed abnormal distribution in cKO spermatids. Rather than outlining axonemal structures as in controls (Fig 5B and C), both proteins accumulated around aggregated centrioles. Moreover, centriole clusters, together with mislocalized sperm tail components, were abundant in the lumen of seminiferous tubules (Fig EV2), suggesting that these aberrant structures are discarded through residual bodies. These findings highlight the critical role of CEP164 in the proper assembly of sperm tail structures.

**Figure 5.**
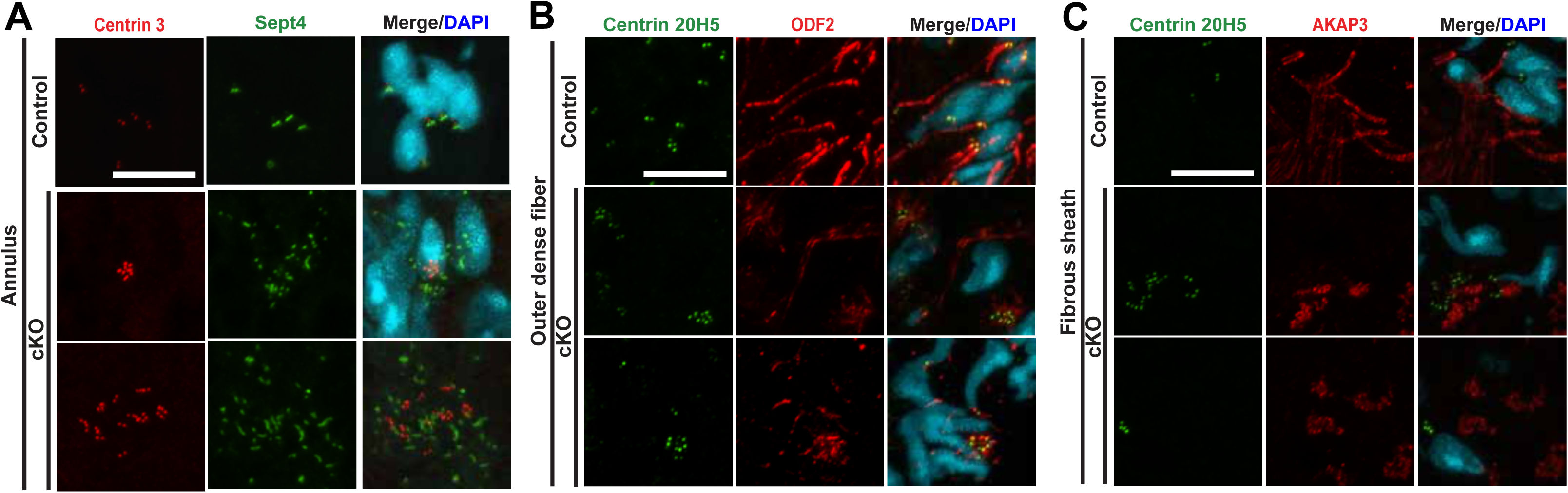
Altered localization of sperm tail structural components in cKO mice. (A-C) IF staining of frozen testis sections using the following antibody combinations: (A) Centrin 3 and Sept4 (annulus component), (B) Centrin [20H5] and ODF2 (outer dense fiber component), and (C) Centrin 20H5 and AKAP3 (fibrous sheath component). Scale bars, 10 µm.

### Centriole clusters first emerge in round spermatids of cKO testes

To determine when centriole clustering first arises in cKO testes, we prepared seminiferous tubule squashes from mice expressing EGFP-Centrin1 and performed IF staining using Centrin 20H5 antibody for precise centriole detection. Quantification across spermatogenesis stages revealed that the majority of pachytene spermatocytes in both control and cKO testes typically contained four centrioles (Fig 6A and B) as reported (Ho *et al*, 2021; Marjanovic *et al*, 2015). In contrast, while most control round spermatids contained two centrioles, 25.2% of cKO round spermatids displayed more than two, and surprisingly, 50.1% lost centrioles entirely (Fig 6C and D). This pattern persisted in elongated spermatids, which showed an increased frequency of cells with supernumerary centrioles (Fig 6E and F). However, the proportion of acentriolar elongated spermatids was markedly reduced, suggesting that round spermatids devoid of centrioles are eliminated during spermatogenesis. Collectively, we conclude that centriole clustering first emerges at the round spermatid stage in the absence of CEP164.

**Figure 6.**
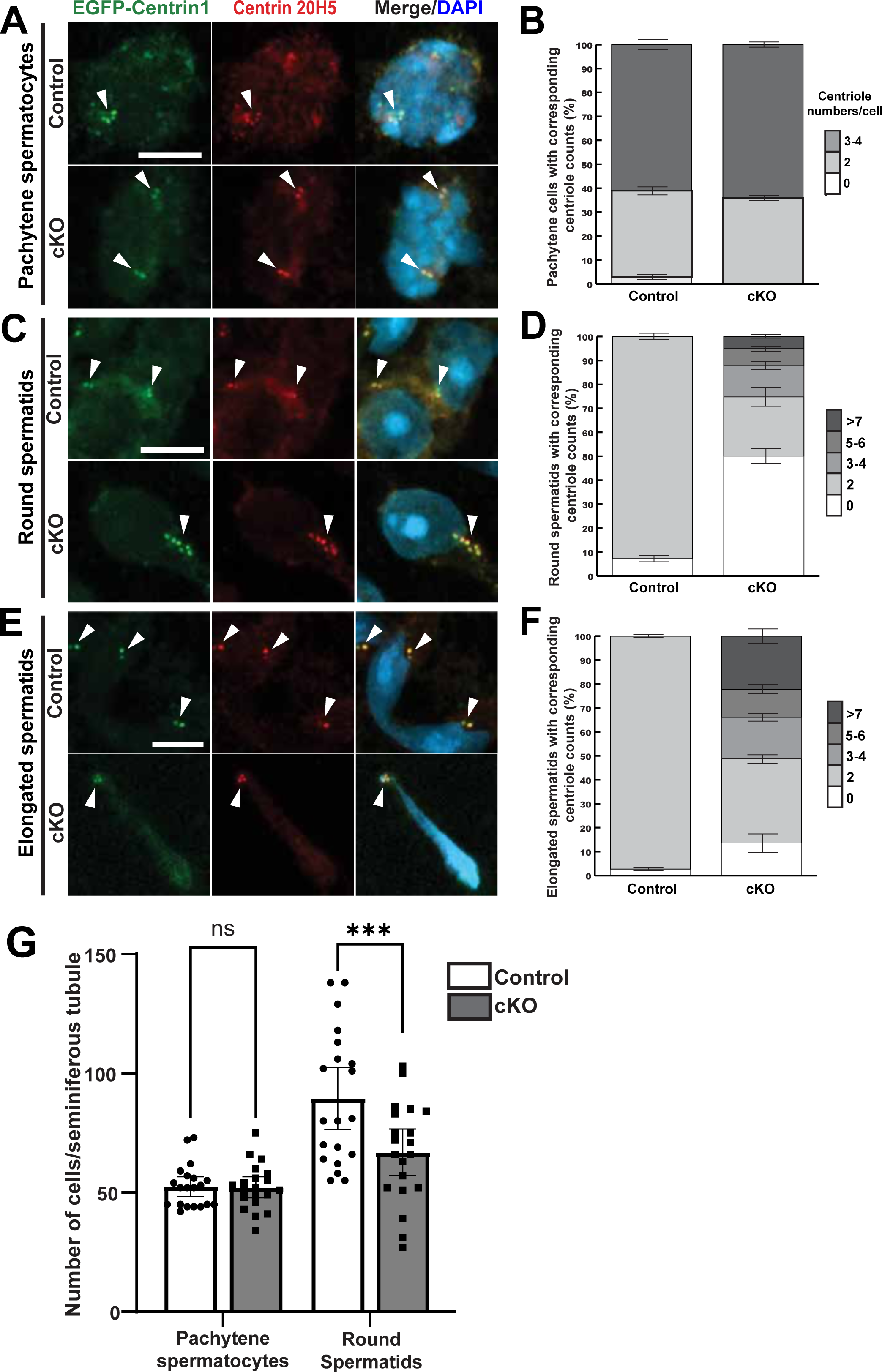
Centriole clusters emerge at the round spermatid stage. (A, C, E) IF staining of seminiferous tubule squashes from mice expressing EGFP- Centrin1 using Centrin 20H5 antibody. Representative images of (A) pachytene spermatocytes, (C) round spermatids, and (E) elongated spermatids are shown. Arrowheads point to centrioles. Scale bars, 5 µm. (B, D, F) Quantification of centriole counts in the corresponding cell types. n = 4 control and n = 3 cKO mice, with 50 pachytene spermatocytes quantified per mouse. n = 6 control and n = 3 cKO mice, with 100 round spermatids quantified per mouse. n = 4 control and n = 3 cKO mice, with 100 elongated spermatids quantified per mouse. Error bars represent ± SEM. (G) Quantification of pachytene spermatocytes and round spermatids in cross-sections of control and cKO juvenile testes (P25-26), immunostained for SYCP1 and SYCP3. n = 2 mice per genotype, with 10 seminiferous tubules analyzed per mouse. Error bars represent ± SEM. Welch’s t test: not significant (ns), ***P < 0.001.

To further substantiate the reduction in round spermatids observed in the cKO testis, we quantified germ cell populations during the synchronized first wave of spermatogenesis by IF staining of P25 and P26 testes for SYCP1 and SYCP3 (Fig EV3). Quantification revealed no significant difference in the number of pachytene spermatocytes between control and cKO testes (Fig 6G). However, the number of round spermatids was significantly reduced in the cKO testis. As no morphological signs of apoptosis were observed, and neither cleaved caspase-3 immunostaining nor TUNEL assays detected apoptotic cells, it is likely that abnormal spermatids in the cKO testis were eliminated through Sertoli cell-mediated phagocytosis (Franca *et al*, 2016; O’Donnell *et al*, 2022; San Agustin *et al*., 2015).

### Centriole pairs are highly mobile and cluster together in cKO round spermatids

We considered two possible mechanisms for the formation of centriole clusters: (1) *de novo* centriole amplification, as reported in cancer cells (Kiermaier *et al*, 2024; Mittal *et al*, 2021) or spermatocytes with dysregulation of polo-like kinases (Skinner *et al*, 2024; Wellard *et al*, 2021), and (2) centriole migration, similar to centrosome migration through nurse cells into oocytes in *D*. *melanogaster* (Bonente *et al*, 2025; Hannaford & Rusan, 2024). To distinguish between these possibilities, we performed live-cell imaging of centrioles in isolated round spermatids from mice expressing EGFP-Centrin1. In control spermatids, centrioles appeared stably anchored to the plasma membrane and exhibited minimal movement (Fig. 7A; Movie EV1). In striking contrast, centrioles in cKO spermatids were highly mobile and dynamic. They frequently associated with one another, likely through intercellular bridges, to form aggregates (Fig. 7A, yellow arrowheads; Movie EV2). No evidence of centriole amplification was detected over three independent 24-hour live-cell recording sessions. These observations suggest that loss of CEP164 and defective basal body docking render centrioles highly mobile, allowing them to pass through intercellular bridges and cluster together, thereby leading to centriole aggregation (Fig. 7B).

**Figure 7.**
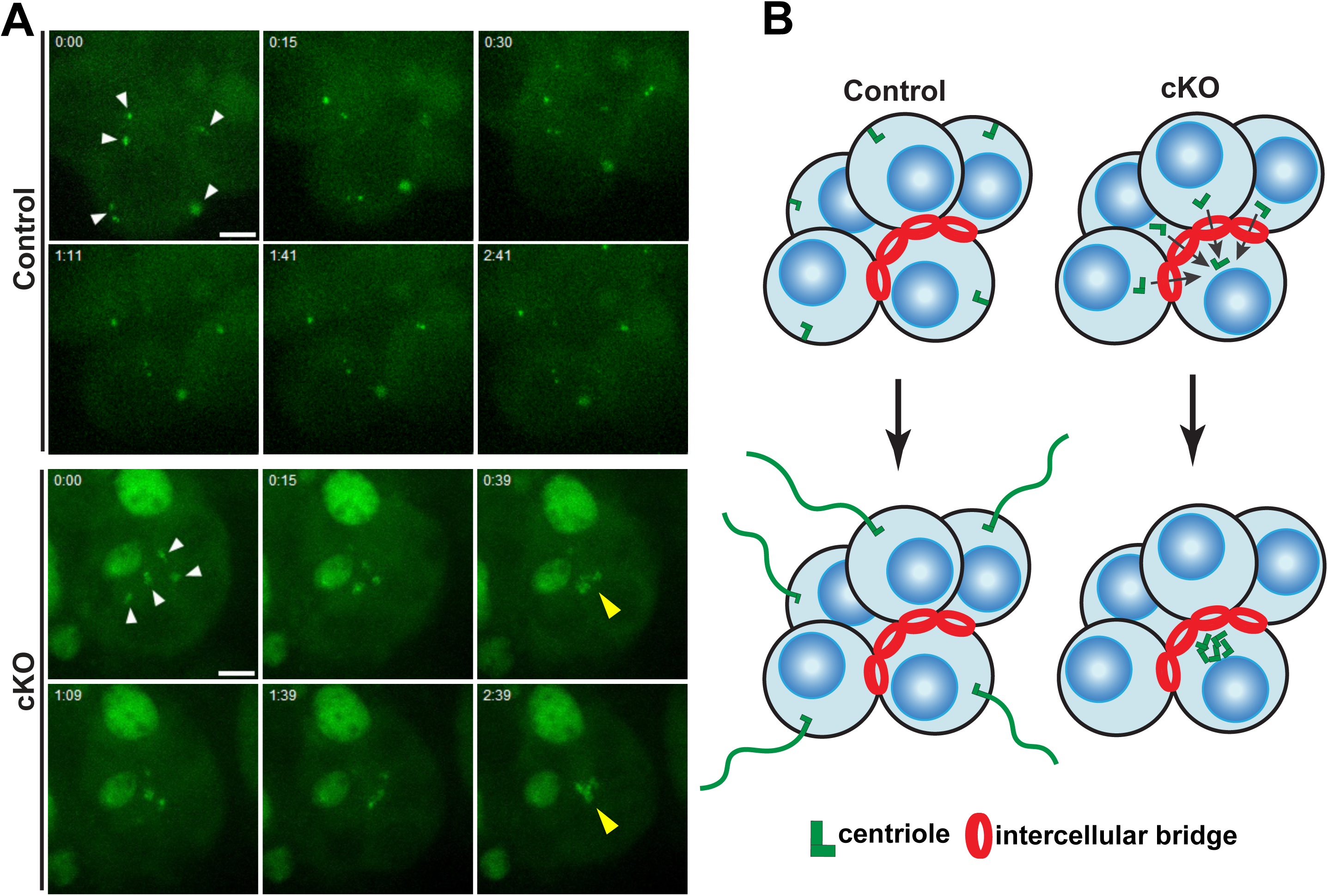
Live-cell imaging of centrioles in round spermatids. (A) Representative spinning disk confocal time-lapse images of EGFP-Centrin1 in round spermatids. Time points are displayed in the upper left corner (h:mm). White arrowheads mark the initial positions of centriole pairs, while yellow arrowheads highlight centriole aggregates in cKO round spermatids. Scale bars, 5 µm. (B) Schematic model illustrating centriole clustering in cKO round spermatids. Defective basal body docking renders centrioles mobile, allowing them to interact via intercellular bridges, ultimately forming large aggregates.

## Discussion

The distal appendage protein CEP164 plays a pivotal role in basal body docking and ciliogenesis in somatic cells. However, its function during germ cell differentiation in mammals has remained largely unknown. In the seminiferous tubules of the adult murine testis, at least two types of cilia have been described: zygotene cilia and sperm flagella (Lopez-Jimenez *et al*., 2022). Although the role of sperm flagella in driving motility and fertilization is well established, the biological significance of zygotene cilia in the testis is not yet understood.

Here, we conditionally ablated CEP164 in male germ cells using the *Stra8*-iCre driver line. In the resulting cKO testis, primary cilia were absent in zygotene spermatocytes (Fig. 2A and C, and Fig. 3A and B). In striking contrast to zebrafish oogenesis, where zygotene cilia are essential for meiosis-specific chromosomal events (Mytlis *et al*., 2022; Mytlis *et al*., 2023), no major lesions were apparent in chromosome pairing and DSB repair in the absence of CEP164 (Fig. 3C-H). These findings suggest that zygotene cilia are dispensable for proper progression of male meiosis in mice. This observation is in agreement with previous reports showing that testis-specific *Kif3A*-KO and *Ift88* hypomorphic mice, deficient in key components of the intraflagellar transport (IFT) machinery essential for ciliogenesis, undergo largely normal spermatogenesis up to the round spermatid stage but subsequently develop pronounced defects in flagellar assembly (Lehti *et al*., 2013; San Agustin *et al*., 2015). However, the status of zygotene cilia and meiotic events were not examined in these studies. Furthermore, a recent study reported that only ∼20% of zygotene spermatocytes possess primary cilia, and that these cilia are not directly associated with bouquet formation during early meiosis (Lopez-Jimenez *et al*., 2022). Therefore, the male infertility observed in cKO mice arises from defects in basal body docking and flagellogenesis during spermiogenesis, rather than abnormalities in meiosis. Interestingly, unlike in mice, germ cell-specific deletion of ciliary genes in zebrafish disrupts meiotic progression by impairing DSB repair and crossover formation, leading to increased germ cell apoptosis and male infertility (Liu *et al*., 2023; Xie *et al*., 2022). This points to an evolutionarily divergent requirement for spermatocyte cilia in regulating gametogenesis across vertebrates.

What are the potential role of zygotene cilia in the mammalian testis? One possibility is that zygotene cilia may function as sensory organelles that regulate spermatogenesis under physiological or environmental stress conditions such as starvation, inflammation, or exposure to chemical or hormonal cues, all of which are known to influence sperm production (Fanjul & Ruiz de Galarreta, 1981; Hasan *et al*, 2022; Sharpe, 2010; Sofikitis *et al*, 2008). Another potential role may involve the blood-testis barrier (BTB), a tight junctional barrier formed by adjacent Sertoli cells that divides seminiferous tubules into the basal compartment and apical (adluminal) compartments. (Luaces *et al*, 2023; Mruk & Cheng, 2015). As preleptotene/leptotene spermatocytes traverse the BTB into a new adluminal environment, zygotene cilia may serve as environmental sensors that monitor local stimuli to modulate germ cell differentiation under certain circumstances. Clearly, further investigation is required to elucidate the precise functions of zygotene cilia during spermatogenesis.

In *D. melanogaster*, the CEP164 ortholog is not essential for basal body docking, flagellogenesis, and male fertility (Hou *et al*., 2023). In contrast, our findings demonstrate that CEP164 is indispensable for male fertility in mice. In the absence of CEP164, spermatogenesis proceeds normally up to the round spermatid stage. However, round spermatids exhibit impaired basal body docking and defective flagellogenesis. Intriguingly, undocked centrioles are highly dynamic and mobile, often associating with one another, likely through intercellular bridges, to form aggregates (Fig 7B). This is accompanied by an increase in acentriolar spermatids, which appear to be eliminated via phagocytosis by Sertoli cells. Centriole clusters are frequently observed in the lumen of seminiferous tubules, suggesting that they are disposed of within residual bodies (Fig. EV2). Centriole clustering in the cKO testis is reminiscent of centriole migration into the oocyte in *D. melanogaster* (Bonente *et al*., 2025; Hannaford & Rusan, 2024). In most metazoans, centrioles are eliminated during oogenesis. In early *Drosophila* oogenesis, a cyst of 16 germ cells interconnected by cytoplasmic bridges forms, and centrioles from the 15 nurse cells migrate through these bridges into the designated oocyte, where they are ultimately discarded as the oocyte matures. Recent evidence indicates that centriole migration through the germline cytoplasm is an evolutionarily conserved process that also occurs in mice (Lei & Spradling, 2016; Niu & Spradling, 2022). An intriguing possibility is that loss of CEP164 and centriolar distal appendages contributes to the migratory behavior of their centrioles in female germ cells. Thus, male germ cells lacking CEP164, with undocked centrioles, appear to adapt a female-like phenotype, characterized by centriole clustering and elimination.

Aberrant centriole clusters are frequently observed in aggressive human cancers and are known to promote multipolar spindle formation, chromosomal instability, and aneuploidy (Kiermaier *et al*., 2024; Mittal *et al*., 2021). The presence of supernumerary centrioles correlates with enhanced metastatic potential and poor clinical prognosis (Kiermaier *et al*., 2024; LoMastro & Holland, 2019). Although centriole amplification is widely regarded as the principal cause of increased centriole numbers, centriole migration through intercellular bridges may present a novel mechanism. The occurrence and function of intercellular bridges in cancer cells have not been thoroughly investigated. However, intercellular bridges or tunneling nanotubes as wide as ∼5 μm in diameter have been reported in cancer cells (Asencio-Barria *et al*, 2019; Vidulescu *et al*, 2004), providing sufficient space for centrioles to travel between connected cells.

In summary, our findings demonstrate that zygotene cilia are dispensable for the proper progression of male meiosis in mice. The distal appendage protein CEP164 is essential for basal body docking, centriole immobilization and retention, and flagellogenesis during murine spermatogenesis. These results illustrate the evolutionarily and biological complexity of gametogenesis, reflecting the presence of species- and sex-specific adaptations in primary cilia function.

## Methods

### Generation of Stra8-iCre;Cep164^fl/fl^ mice

All animal procedures were conducted in accordance with NIH guidelines and approved by the Institutional Animal Care and Use Committee (IACUC) of Stony Brook University. The generation and genotyping of *Cep164^fl/fl^* mice have been described previously (Siller *et al*., 2017). *Stra8*-iCre mice were obtained from the Jackson Laboratory (RRID:IMSR_JAX:008208). Mice homozygous for the *Cep164* flox allele and hemizygous for the *Stra8-iCre* transgene (*Stra8-iCre;Cep164^fl/fl^*, cKO) mice were generated by breeding *Stra8-iCre;Cep164^fl/+^* mice with *Cep164^fl/fl^* mice. Adult mice (2-5 months of age) were used for all the experiments unless otherwise specified. *Cep164^fl/fl^* mice were used as controls throughout the study, except for Fig 6G, in which P25 *Stra8- iCre;Cep164^fl/+^* and P26 *Cep164^fl/+^* littermates were used as controls. Animals were maintained under pathogen-free conditions on a 12:12 hour light:dark cycle with free access to food and water in the Division of Laboratory Animal Resources at Stony Brook University.

### Quantification of cauda epididymal sperm

Cauda epididymal sperm were isolated and counted as previously described (Hoque *et al*., 2024). Briefly, 2-5-month-old mice were euthanized, and a single cauda epididymis was dissected and minced in a 35-mm petri dish containing 1 ml PBS (pH7.4). The tissue was incubated at 37°C for 15 minutes to allow sperm release. The suspension was collected, centrifuged, and the sperm-containing pellet was resuspended in 1 ml PBS. Sperm were counted using a hemocytometer with appropriate dilutions.

### Histological staining

Testes and epididymides were collected from 2-3-month-old mice, fixed overnight in Bouin’s solution (Ricca Chemical), and embedded in paraffin. Sections of 5-10 µm were cut, stained with hematoxylin and eosin (H&E; Poly Scientific R&D Corp.) or Periodic Acid-Schiff Kit (PAS; Sigma-Aldrich), and mounted with Permount (Fischer Scientific).

### Immunofluorescence staining

For immunofluorescence (IF) staining of paraffin-embedded tissues, sections were deparaffinized, rehydrated, and subjected to antigen retrieval in sodium citrate buffer (10 mM sodium citrate, 0.05% Tween 20, pH 6.0) using an autoclave for 17.5 minutes.

Sections were permeabilized with 0.5% Triton X-100 in PBS for 5 minutes, washed three times with 0.05% Tween 20 in PBS (5 minutes each), and blocked for 1 hour at room temperature in 5% goat serum prepared in antibody dilution solution (2% BSA in PBS). Primary antibodies were applied in antibody dilution buffer and incubated overnight at 4°C. After three washes with 0.05% Tween 20 in PBS, sections were incubated with secondary antibodies for 1 hour at room temperature, washed three additional times with 0.05% Tween 20 in PBS, counterstained with DAPI for 2 minutes, and mounted with Fluoromount-G (Southern Biotech).

For IF staining of frozen tissues, freshly isolated testes were embeded in OCT compound (Tissue Tek), frozen at -80°C overnight, and sectioned at 10-µm thickness. Sections were air-dried for 15 minutes and fixed in cold methanol:acetone (1:1) for 10 minutes. Subsequent steps were performed as described above, except the Triton X- 100 permeabilization step was omitted. Primary and secondary antibodies are listed in Table EV1.

### Seminiferous tubule squash and meiotic chromosome spread preparations

Seminiferous tubule squashes were performed as previously described(Wellard *et al*, 2018) with modifications. Testes from 2-4-month-old mice were harvested, detunicated, and individual seminiferous tubules were incubated in fix/lysis solution (0.8% PFA and 0.1% Triton X-100 in PBS) for 5 minutes. Seminiferous tubules were then cut into 5-20 mm fragments and transferred onto charged slides pre-coated with fix/lysis solution. Coverslips were placed over the tubules, and pressure was applied with the heel of the palm for 30 seconds. Slides were immediately flash-frozen in liquid nitrogen for ∼20 seconds and used immediately for IF staining or stored at -80 °C. For IF staining, slides were reimmersed in liquid nitrogen and coverslips were removed with a razor blade.

Meiotic chromosome spreads were prepared as previously described (Dia *et al*, 2017) with modifications. Testes from 2-3-month-old mice were harvested, detunicated, and incubated in hypotonic extraction buffer (HEB; 30 mM Tris-HCl pH8.2, 50 mM sucrose, 17 mM trisodium citrate dihydrate, 5 mM EDTA, 0.5 mM DTT, and 0.1 mM PMSF) for 1 hour. Testis clumps (∼3 mm in diameter) were then minced in 100 mM sucrose solution and spread onto slides pre-coated with fixative solution (1% PFA, 0.15% Triton X-100). Slides were dried in a covered humidified chamber for 3 hours, followed by 30 minutes in an open humidified chamber. After three 5-minute PBS washes, slides were air-dried vertically for 20 minutes and either used immediately for IF staining or stored at -80°C.

### Transmission electron microscopy

TEM was performed as described previously (Hoque *et al*., 2024; Siller *et al*., 2017). Testes from 2-month-old mice were dissected, cut into ∼1 mm^3^ pieces, and fixed overnight at 4°C in 4% PFA and 2% glutaraldehyde in PBS. Samples were rinsed several times with PBS, post-fixed with 2% osmium tetroxide for 1 hour, dehydrated through a graded ethanol series, followed by acetonitrile, and embedded in Embed 812 resin (Electron Microscopy Sciences) for 48 hours at 60°C. Ultrathin sections (60-90 nm) were cut on a Leica EM UC7 ultramicrotome, collected on Formvar-coated slot copper grids, and counterstained with 1% methanolic uranyl acetate and 0.3% aqueous lead citrate. Images were acquired on a FEI Tecnai G2 12 BioTwin TEM equipped with an XR-60 CCD digital camera system (Advanced Microscopy Techniques).

### Live-cell imaging of centriole dynamics in spermatids

Testes were collected from 2-3-month-old mice expressing EGFP-Centrin1, detunicated, and seminiferous tubules were transferred to a sterile Petri dish containing PBS. Tubule segments at spermatogenic stages I-V, which contain round spermatids, were identified by phase-contrast microscopy using the transillumination method (Kotaja *et al*, 2004). Selected segments were isolated and transferred to 35-mm culture dishes containing MEMα (Gibco) supplemented with 10% KnockOut Serum Replacement (KSR; Gibco) and 100 U/mL penicillin-streptomycin. Tubules were gently minced to release clumps of spermatids, which are then allowed to disperse. Live-cell imaging was performed for 24 hours at 15-minute intervals using a Nikon Eclipse Ti2-E CrestOptics X-Light V3 spinning disk confocal system equipped with a 40×/0.90 NA objective and an environmental chamber at 37°C and 5% CO2.

### Image acquisition and analysis

For histological staining, images were acquired using a Leica DMI6000B microscope equipped with either a DFC7000T camera and either a 20×/0.50 NA or 40×/0.75 NA objective. IF images were obtained using a Leica SP8X confocal microscope with a HCPL APO 100×/1.40 NA oil-immersion objective or a Nikon AXR confocal microscope with either a 20x/0.80 NA or 60x/1.42 NA oil-immersion objective in Galvano scan mode. Image analysis was performed using Leica Application Suite X or Nikon NIS-Elements software. Final images were generated using ImageJ, Adobe Photoshop, and Adobe Illustrator.

### Statistical analysis

Two-tailed Welch’s t tests were used to determine statistical significance in all cases. P < 0.05 was considered to be significant. Asterisks were used to indicate P values as follows: **P < 0.01; ***P < 0.001; and ns, not significant.

## Acknowledgements

We would like to thank Dr. Mohammed Hoque for his assistance with the initial experiments. We acknowledge the Central Microscopy Imaging Center Core at Stony Brook University for assistance with confocal microscopy and TEM and the Research Histology Core Laboratory for assistance with histological preparations. This work was supported by grants from the National Institute of Child Health and Human Development (R01HD109232 to K.-I. Takemaru and R01HD109232-S1 to J.J. Chen) and from the National Institute of General Medical Sciences (R35GM142875 to M. Mak).

## EV Figure Legends

**Figure EV1.**
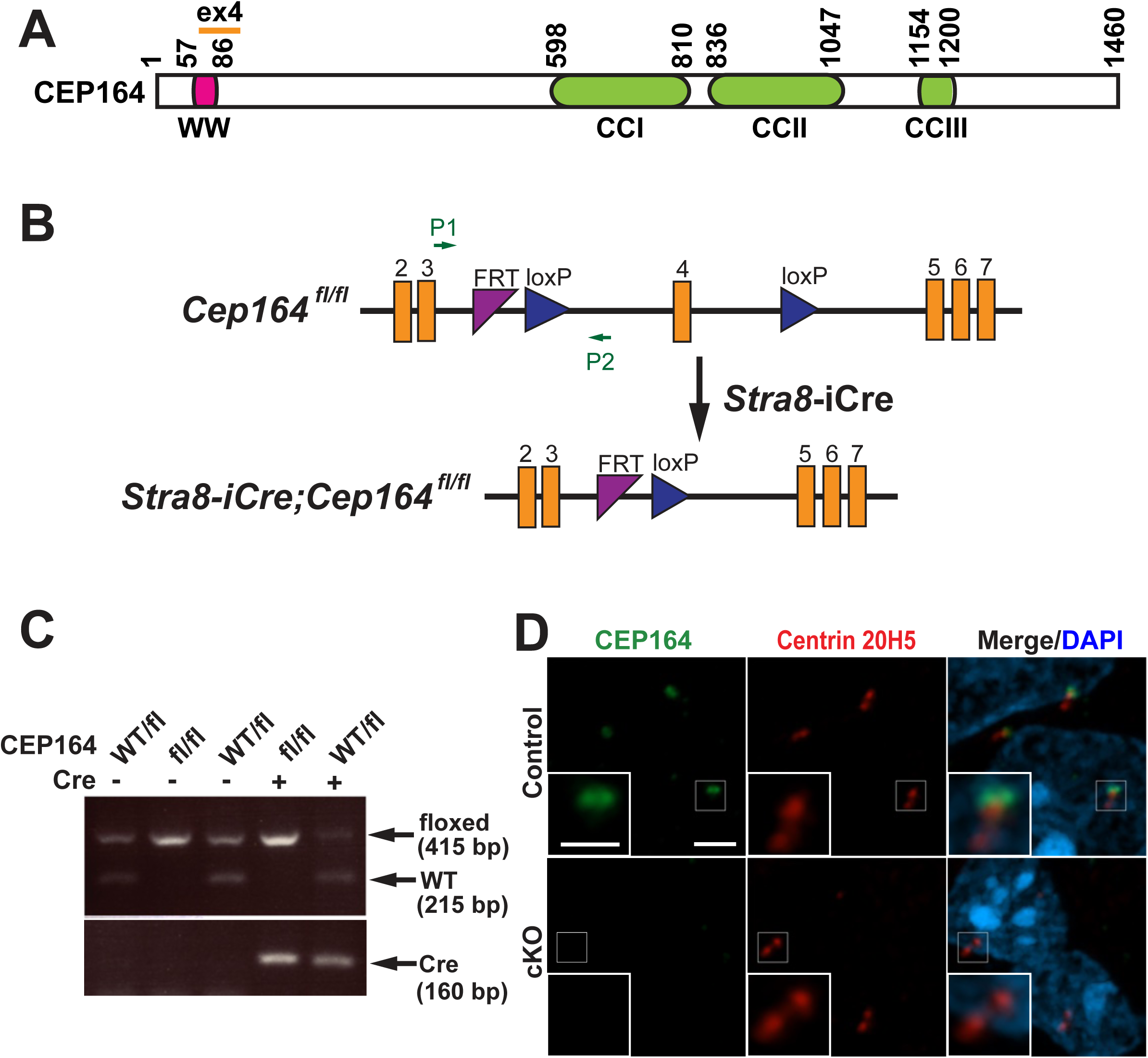
Generation of *Stra8-iCre;Cep164^fl/fl^* mice. (A) Schematic of the CEP164 protein domain organization, showing a WW domain and three coiled-coil motifs (CCI-CCIII). (B) Strategy for generating *Stra8-iCre;Cep164^fl/fl^* mice. The locations of genotyping primers are indicated by green arrows (P1 and P2). (C) PCR-based mouse genotyping. (D) IF staining of adult frozen testis sections using CEP164 and Centrin 20H5 antibodies. Scale bars, 2 and 1 µm (insets).

**Figure EV2.**
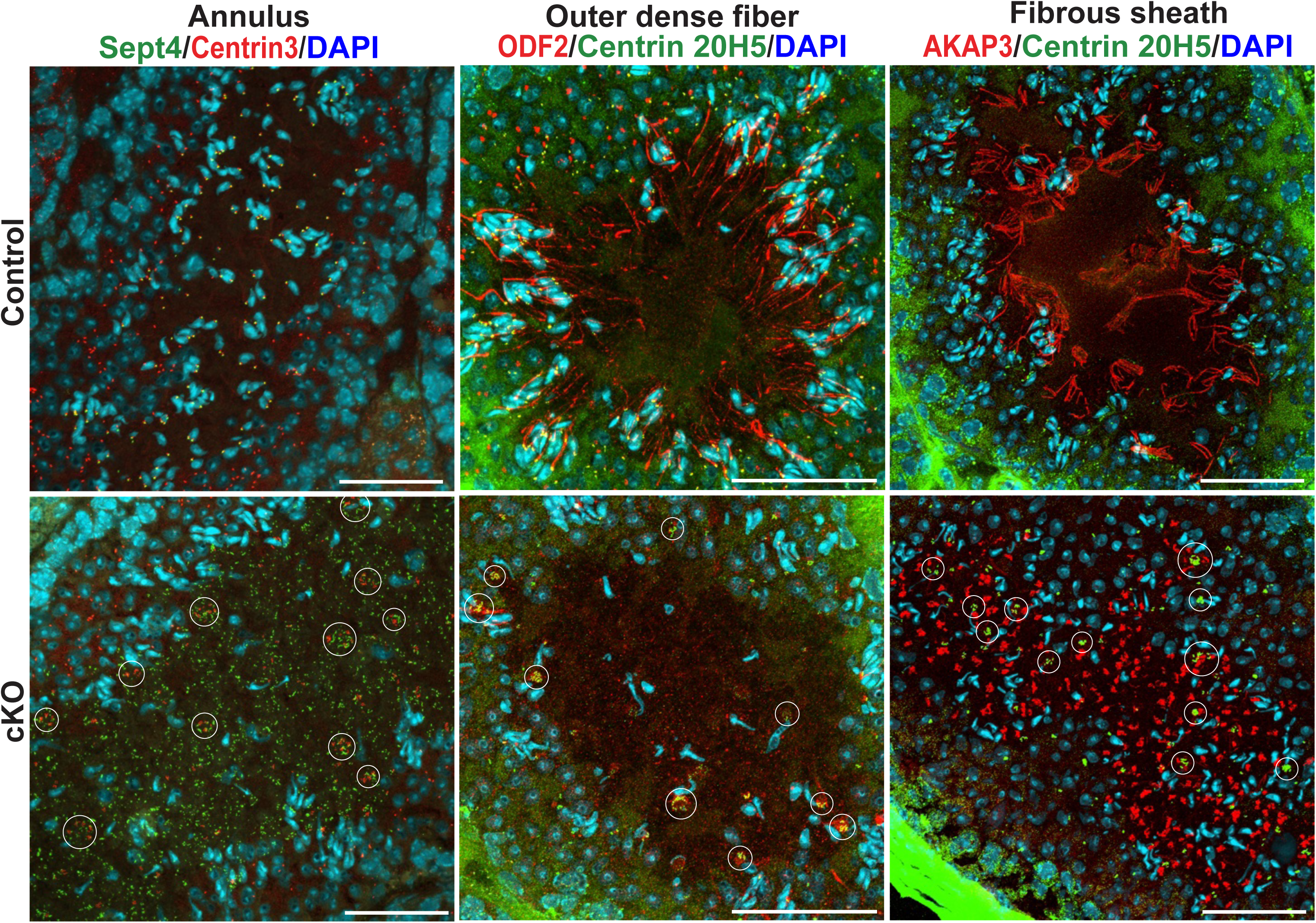
IF analysis of sperm tail structural components in adult testes. Frozen testis sections were immunostained using the following antibody combinations: Centrin 3 and Sept4 (annulus component), Centrin [20H5] and ODF2 (outer dense fiber component), and Centrin 20H5 and AKAP3 (fibrous sheath component) as indicated. Centriole clusters are outlined by circles. Scale bars, 50 µm.

**Figure EV3.**
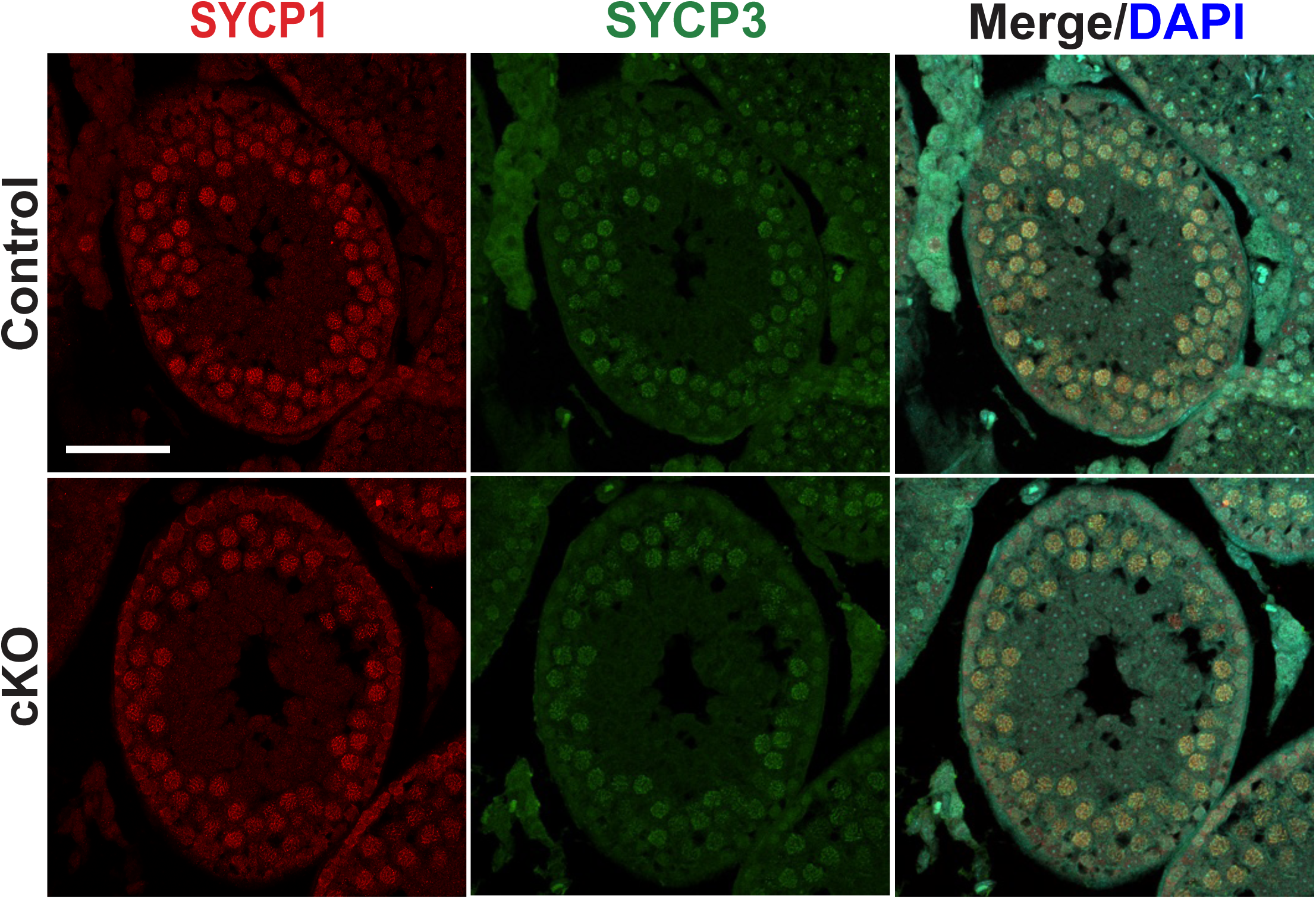
Representative IF images of P25 testis sections immunostained for SYCP1 and SYCP3. Paraffin testis sections from P25 littermates during the synchronized first wave of spermatogenesis were immunostained for SYCP1 and SYCP3. Scale bars, 50 µm.

**Movie EV1. Live-cell time-lapse imaging of centrioles in control spermatids.** Control spermatids were imaged at 15-minute intervals. Arrowheads indicate the initial positions of centriole pairs. Scale bar, 5 µm.

**Movie EV2. Live-cell time-lapse imaging of centrioles in cKO spermatids.**

cKO spermatids were imaged at 15-minute intervals. Arrowheads indicate the initial positions of centriole pairs. Floating cells exhibiting intense green fluorescence are seen. Scale bar, 5 µm.

**Table EV1.** Antibodies used for immunofluorescence staining.

| Name | Species | Manufacturer | Catalog number | RRID |
| --- | --- | --- | --- | --- |
| Acetylated α-tubulin [6-11B-1] | Mouse | Sigma-Aldrich | T6793 | AB_477585 |
| AKAP3 | Rabbit | Proteintech | 13907-1-AP | AB_2273887 |
| Centrin [20H5] | Mouse | Sigma-Aldrich | 04-1624 | AB_10563501 |
| Centrin 3 [3E6] | Mouse | Abnova | H00001070-M01 | AB_537701 |
| CEP164 [17] | Mouse | Sigma-Aldrich | SAB2702371 | N/A |
| ODF2 | Rabbit | Proteintech | 12058-1-AP | AB_2156630 |
| Phospho-H2A.X (Ser139) | Rabbit | Sigma-Aldrich | 07-164 | RRID:AB_11213838 |
| Rad51 | Rabbit | Sigma-Aldrich | PC130 | AB_2238184 |
| SEPT4 | Rabbit | Proteintech | 12476-1-AP | AB_2187151 |
| SYCP1 | Rabbit | Invitrogen | PA1-46254 | AB_2302734 |
| SYCP3 [D-1] | Mouse | Santa Cruz | sc-74569 | AB_2197353 |
| Anti-Mouse IgG Alexa Fluor 488 PLUS | Goat | Invitrogen | A32723 | AB_2633275 |
| Anti-Mouse IgG1 Alexa Fluor 488 | Goat | Invitrogen | A21121 | AB_2535764 |
| Anti-Mouse IgG2a DyLight 488 | Goat | Jackson ImmunoResearch Labs | 115-485-206 | N/A |
| Anti-Mouse Alexa Fluor 555 PLUS | Goat | Invitrogen | A32727 | AB_2633276 |
| Anti-Mouse<br>Alexa Fluor<br>594 PLUS | Goat | Invitrogen | A32742 | AB_2762825 |
| Anti-Mouse<br>IgG2a<br>DyLight 549 | Goat | Jackson<br>ImmunoResearch<br>Labs | 115-505-206 | N/A |
| Anti-Mouse<br>IgG2b<br>DyLight549 | Goat | Jackson<br>ImmunoResearch<br>Labs | 115-505-207<br>(91425) | N/A |
| Anti-Mouse<br>IgG1 DyLight<br>649 | Goat | Jackson<br>ImmunoResearch<br>Labs | 115-495-205 | N/A |
| Anti-Rabbit<br>Alexa Fluor<br>488 PLUS | Goat | Invitrogen | A32731 | AB_2633280 |
| Anti-Rabbit<br>Alexa Fluor<br>555 PLUS | Goat | Invitrogen | A32732 | AB_2633281 |
| Anti-Rabbit<br>Alexa Fluor<br>594 PLUS | Goat | Invitrogen | A32740 | AB_2762824 |
| Peanut<br>Agglutinin<br>(PNA),<br>Rhodamine |  | Vector Labs | RL-1072-5 | N/A |

## Notes

### Competing Interest Statement

The authors have declared no competing interest.

